# Antagonizing the corticotropin releasing hormone receptor 1 with antalarmin reduces the progression of endometriosis

**DOI:** 10.1101/317867

**Authors:** Annelyn Torres-Reverón, Leslie L. Rivera-Lopez, Idhaliz Flores, Caroline B. Appleyard

**Author notes:** **Corresponding Author:** Annelyn Torres-Reveron, PhD, University of Texas at Rio Grande Valley, School of Medicine, 1214 W Schunior Street, Edinburg TX 78541, E.

## Abstract

Endometriosis is a disorder in which endometrial tissue is found outside the uterus causing pain, infertility and stress. Finding an effective and long-term treatment for endometriosis still remains one of the most significant challenges in the field. Corticotropin releasing hormone (CRH) is one of the main signaling peptides within the hypothalamic pituitary adrenal (HPA) axis released in response to stress. CRH can affect nervous and visceral tissues such as the uterus and gut via activation of two types of CRH receptors: CRHR1 and CRHR2. Our aim was to determine if blocking CRHR1 with antalarmin will reduce endometriosis progression. First, we induced endometriosis in female rats by suturing uterine horn tissue next to the intestinal mesentery and allowed to progress for 7 days. We determined that after 7 days, there was a significant increase in CRHR1 within endometriotic vesicles as compared to normal uterus. A second group of rats received endometriosis but also antalarmin (20 mg/kg, i.p.) during the first 7 days after surgery. As separate group of sham surgery rats served as controls. Endometriosis was allowed to progress until 60 days after surgery. At time of sacrifice, rats were tested for anxiety behaviors and endometriotic vesicles, and uterus were collected. Rats with endometriosis that received antalarmin significantly reduced the size (67% decrease) and number (30% decrease) of endometriotic vesicles. Antalarmin also prevented the increase in CRH and CRHR1 within endometriotic vesicles but not of glucocorticoid receptor. Behaviorally, endometriosis increased anxiety in the zero-maze test but antalarmin did not modify it. Our data provides the first demonstration for the effective use on CRHR1 antagonist for the treatment of endometriosis with promising effects for long-term therapy of this debilitating disease.

## Introduction

Corticotropin releasing hormone (CRH) is one of the main signaling molecules of the hypothalamic pituitary adrenal (HPA) axis. CRH has a myriad of physiological effects that include behavioral, endocrine, autonomic and immune responses (1,2). CRH acts mainly by binding to CRH receptors type 1 (CRHR1) and type 2 (CRHR2) with a 10-fold affinity for the CRHR1 versus CRHR2 (3). CRH receptors belong to the superfamily of G-protein coupled receptors and typically effect cellular activity via coupling to adenylate cyclase (3). CRHR1 is abundant in the brain (4) as well as in adrenal glands, uterine and colonic tissues, and lymphocytes, among others (5–7). Eleven splice variants of the CRHR1 receptor have been identified (8), with a tissue specific expression pattern (9,10). In addition, the CRH paralog, urocortin 1 (UCN1) can bind and activate both the CRHR1 and R2 (11).

Due to the variety of physiological activities that the CRH system exerts, CRHR1 antagonists have been clinically used for more than three decades for a variety of conditions. For example, CRHR1 antagonists have been tested for the treatment of disorders including depression (12), irritable bowel syndrome(13) (IBS), and proposed as a possible treatment for anxiety disorders (14). In fact, phase II/III clinical trials are undergoing or have been completed for depression, IBS and anxiety (15). Antalarmin is a CRHR1 antagonist that has been widely used in animal research to investigate CRH effects on reproduction, inflammation, addictive disorders, sleep disorders, among others (16). Antalarmin is a non-peptide molecule that readily crosses the blood-brain-barrier. Both, anti-stress and anti-inflammatory activities of antalarmin have been documented in animal studies (17).

Endometriosis is a chronic inflammatory disorder defined as the presence of endometrial-like tissue (e.g., glands and stroma) outside the endometrial cavity. This condition is characterized by peritoneal inflammation resulting in severe and chronic pelvic pain, and often infertility (18). Endometriosis can be commonly misdiagnosed as irritable bowel syndrome (IBS)(19) due to overlap in common symptoms and perhaps mechanisms of disease progression involving aberrant activation of inflammatory cascades. The causes of endometriosis onset are unknown; however, a relationship between stress, hypothalamic pituitary adrenal axis (HPA) dysregulation, and endometriosis severity has been documented by others and our own work in the rat model of endometriosis (20–23). Strong evidence (from both human and animal studies) suggest that abnormal functioning of the HPA axis, release of CRH and/or the inflammatory response system disrupts feedback of both neuroendocrine and immune systems contributing to the development of the disease (24,25). CRH and CRH receptors are abundant in female reproductive tissues and this axis has been shown to regulate several reproductive functions (26,27), mostly mediating pro-inflammatory activities such as ovulation, luteolysis and blastocyst implantation (2). Despite the well-documented role of CRH receptor in stress related disorders, reproductive function and inflammation, no previous study has addressed the potential role of CRHR1 blockade in the treatment of endometriosis.

In the current study, we took advantage of the well-established auto transplantation rat model of endometriosis to investigate the effects of the CRHR1 receptor antagonist antalarmin in endometriosis. Given the role of CRHR1 in inflammation, we first tested whether this receptor was up regulated in ectopically implanted endometrium shortly after disease induction. Following this, we administered antalarmin, early during endometriosis establishment to test whether it could block vesicle formation in this model. We hypothesized that blockade of CRHR1 during the first week after endometriosis induction will reduce the initiation of inflammatory processes that lead to endometriosis vesicle establishment and subsequent development. In addition, we hypothesized that CRHR1 blockade at central levels may reduce stress associated behaviors previously linked to endometriosis such as anxiety and depression. Data presented herein suggest that the CRHR1 antagonist antalarmin might function as a completely new line of treatment for women suffering from endometriosis, highly needed in the management of this debilitating and still incurable disease.

## Materials and Methods

### Animals and experimental groups

Female Sprague Dawley rats of 60 days old were used in the experiments (weighing between 190 – 220 grams). Rats were housed two per cage and kept in a 12-hour light/dark cycle with food and water ad libitum. All experimental procedures were approved by the Ponce Health Sciences University and the University of Texas at Rio Grande Valley Institutional Animal Care and Use Committees and adhere to the NIH Guide for the Care and Use of Laboratory Animals. Rats were weighed twice per week to monitor their adequate development and once a week during drug administration period. Group 1 consisted of 32 female rats that underwent endometriosis induction or sham surgery (described below) and were sacrificed 7 days after surgery (surgery - Day 0; Figure 1). This experiment was carried out to quantify the levels of CRHR1 receptors at 7 days after surgery and thus assess the feasibility of using a CRHR1 antagonist during this period. Group 2 consisted of 40 female rats that underwent endometriosis induction or sham surgery and received the CRHR1 antagonist antalarmin or vehicle control from days 1-7 after surgery. Parallel to this group, a separate group of 11 rats underwent sham surgery and were left untreated until day 60 after surgery. Only rats with regular estrous cycles were used in the experiments as assessed by vaginal smear lavage during the 7 days prior to surgery and 7 days before sacrifice.

**Figure 1:**
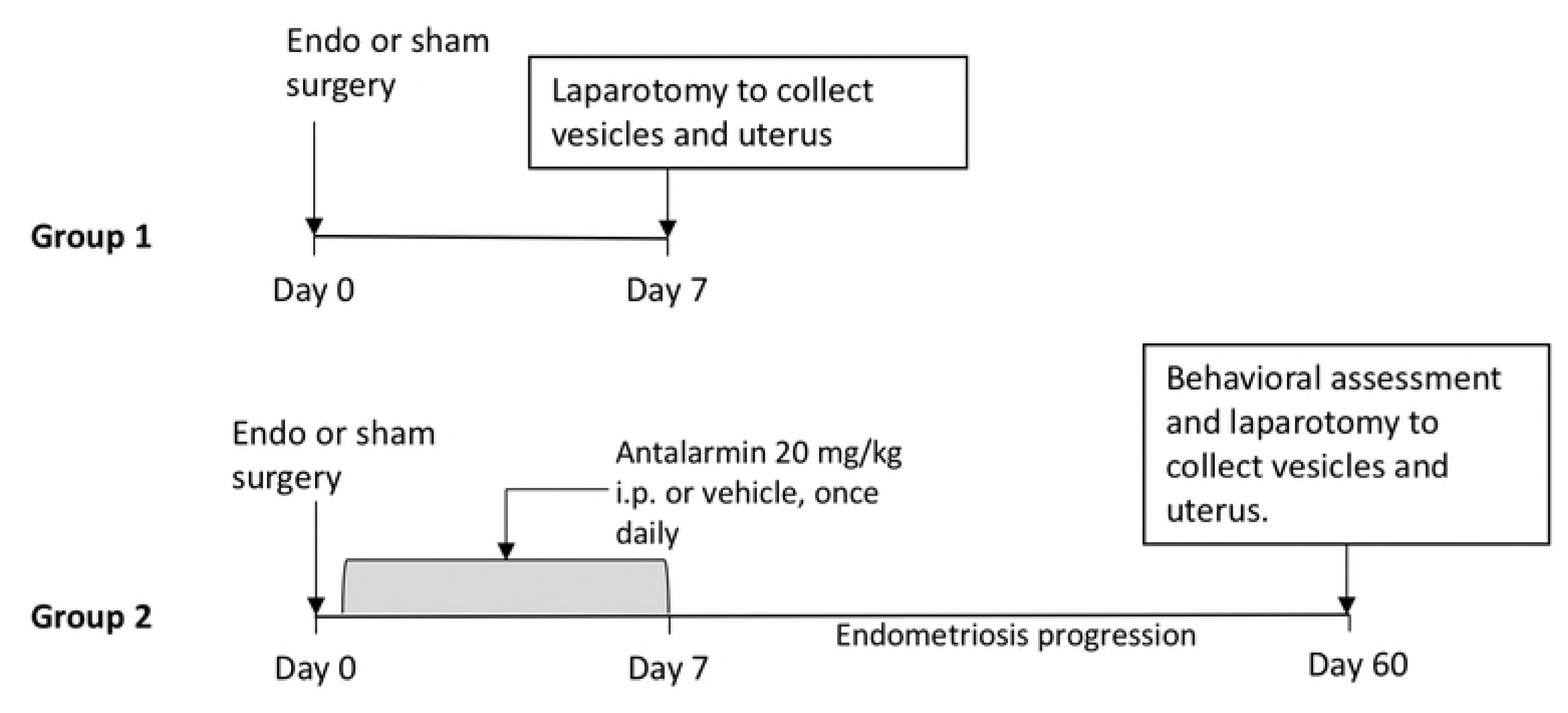
Diagram of experimental protocols. Rats in Group 1 received endometriosis or sham surgery and were allowed to progress for 7 days. Rats in Group 2 received sham surgery or endometriosis. Rats from the endometriosis group were injected with either vehicle or antalarmin for seven consecutive days after surgery and allowed to progress for 53 additional days. During the endometriosis progression period, animals were undisturbed except for weekly weighing done at the same time of cage changing.

### Drug administration

Animals in Group 2 received 1 daily injection (between 09:00 – 10:00 hours) for 7 consecutive days of antalarmin (N-butyl-N-ethyl-[2,5,6-trimethyl-7-(2,4,6-trimethylphenyl)-7H-pyrrolo[2,3-d]pyrimidin4-yl]-amine; Tocris Bioscience, Bristol, UK) suspended in a vehicle composed of 10% Tween 80 and distilled water and given intraperitoneally at 20 mg/kg in a volume of 1 ml/kg. This dose of antalarmin was chosen based on previous published work from Cippitelli, et al., (2012) (28) dose-response study showing that the 20 mg/kg i.p. administration was the most effective dose to block withdrawal behaviors, thus it readily entered the blood-brain barrier in addition to the peripheral tissues. Antalarmin injections started the morning following surgery. After day 7, rats were left undisturbed except for cage change and weighing twice a week.

### Endometriosis induction

Endometriosis was surgically induced as previously described elsewhere (29,30). Briefly, rats were anesthetized with isoflurane and four pieces of the right uterine horn were auto transplanted to 4 different blood vessels in the intestinal mesentery. The control group were sham operated animals for which the right uterine horn was massaged for 2 minutes and sutures were placed in the intestinal mesenteric area with no uterine implants. For group 1, sham or endometriosis operated rats were allowed to progress for 7 days after surgery. For group 2, endometriosis was allowed to progress for 60 days before sacrificing, similar to our previous report (31;30).

### Behavioral assessment

One day before behavioral assessment and sacrifice a subset of rats (6 vehicle and 6 antalarmin) were subjected to an acute episode of swim stress of 10 min and compared to no stress controls to assess how rats respond to acute activation of the HPA axis. For this, animals were placed in a Plexiglas tank for 10 min in water at 25°C (modified from (32)). Rats were towel dried and kept in warm cage after swim until fur dried. The next day, we used two behavioral tasks to assess anxiety behaviors. The open field test is used to quantify exploratory and locomotor activity of a rodent in an open arena. The apparatus used was a square wood arena (91 x 91 x 38 cm) with overhead light illumination and video monitoring to record animal activity using Any-Maze software (Stoelting, Wood Dale, Illinois). We quantified the following behaviors during 20 minutes: 1) total distance moved, 2) time spent moving, 3) time spent in the center of the arena, 4) time spent near the walls of the arena (defined by the 15 cm of floor arena closest to the walls) and 5) total fecal pellets. The more time the animal spends in the center of the arena compared to the space adjacent to the wall is considered as having less anxiety. At the end of the testing period, animals were returned to the home cage and after a 5-min break were tested in the elevated zero-maze.

The elevated zero-maze is very similar to the more traditional elevated plus maze test, with the advantage of not having a neutral (undefined) zone in the middle. The apparatus consisted of a circle with an arm width of 10cm and elevated 40cm from floor. Two sections of the circle were open without walls and two enclosed by 40 cm high walls. Rats were placed in the intersection of an open arm, facing the closed arm and opposite to the experimenter. Rats were allowed to run the maze for 5 consecutive minutes and recorded using the Any-Maze software (Stoelting, Wood Dale, Illinois). The following parameters were analyzed by the Any-Maze program: 1) total distance travelled in the maze, 2) time spent in the open/closed arms and 2) number of entries made by the rodent onto the open/closed arms. When 60% of the animal body entered the arm, the program counted it as an entry. In addition, we quantified total fecal pellets in the maze. The more time the animal spends in the open arms is considered as having less anxiety. After the 5-min testing period, the rat was returned to the home cage and immediately anesthetized with an overdose of 65% sodium pentobarbital to proceed with laparotomy. The maze was thoroughly cleaned with 70% alcohol solution and allowed to dry before testing the next rat.

### Sample Collection and Processing

We verified that the animals were deeply anesthetized. Rats were weighed, and a cytological smear taken to verify stage of the estrous cycle. The peritoneal and thoracic cavities were opened, and a blood sample was collected directly from the heart. Following this, we collected peritoneal fluid using a sterile plastic pipette. Then, we examined for the presence of endometriosis vesicles. The implants that developed into vesicles were excised from the mesentery, weighed and measured using a digital caliper. Classification of vesicles was carried out as previously described (23,33) and assigned the following grades: grade 1= disappeared; grade 2= 0.01- 1.99 mm; grade 3= 2 - 4.49 mm; grade 4= 4.5 – 5.99 mm; grade 5= 6.0 mm or larger. In sham animals, we counted and collected the empty suture sites. In addition to the endometriosis vesicles, we collected the adrenal glands, removed all surrounding fatty tissue and weighed them. We also collected tissues from colon, and the left uterine horn. All tissues were flash frozen and stored at -80° until further processing.

### Enzyme linked immunosorbent assays (ELISA)

Serum and peritoneal fluid samples from animals were tested for levels of corticosterone, adrenocorticotropic hormone (ACTH) and the pro-inflammatory cytokine IL-6 following instructions in the commercial kits. The following kits were used: Corticosterone rat/mouse kit (Cat. # 79175; IBL America, Minneapolis, MN); Rat IL-6 pre-coated ELISA kit (Cat. #437107; BioLegend, San Diego, CA); Mouse/rat ACTH ELISA kit (Cat. #AC018T-100, Calbiotech, El Cajon, CA).

### RNA isolation and cDNA synthesis

Endometriosis vesicles and normal uterine tissue, from endometriosis or sham rats were lysed in an RLT buffer (Qiagen, Germantown, MD) using the Bullet Blender Tissue Homogenizer (Next Advance, Averill Park, NY). The total RNA from the lysates were extracted according to RNAeasy Mini Kit manufacturer’s protocol (Qiagen, Germantown, MD). RNA concentration and purity were measured on a NanoDrop 2000 UV spectrophotometer (Thermo Scientific, Wilmington, USA). Concentration and quality of RNA samples were acquired based on the ratio of absorbance at 260/280 nm in the spectrophotometer. To carry out the synthesis of cDNA from RNA samples a total reaction volume of 20 μl including 0.1μg of total RNA concentration and synthesis reagents was used. We used the iScript cDNA Synthesis Kit according to manufacturer’s protocol (Bio-Rad, Hercules, CA). Reactions were carried out in T-100 thermal cycler (Bio Rad, Hercules, CA). RT-PCR running method was as follows: 25°C for 5 min, 42°C for 30 min, 85°C for 5 min. Samples were stored at -80° C for later experimentation or qRT-PCR.

### Quantitative real time PCR protocol

We used real time quantitative PCR (qPCR) to evaluate changes in mRNA expression. For this, we used 25 μl of a total volume of reaction assay with 1:10 dilution of cDNA with IQ SyBR Green Supermix (Bio Rad Hercules, CA) in a 96 well plate according to the manufacturer protocol and amplified in a Quant Studio 12K Flex Real time PCR System (Applied Biosystems, Carlsbad, CA). Commercial primers for CRH, UCN1 CRHR1, CRHR2 and GR were purchased from Qiagen (Germantown, MD). Real time PCR cycles protocol was as follows: 95°C for 10 min. for enzyme activation followed by 40 cycles of denaturing at 95 °C for 15 sec. and annealing at 60°C for 1 min. All changes in gene expression were normalized against GAPDH of each sample. CT values and changes per gene expression level were automatically analyzed by the Quantstudio 12K Flex Software (Applied Biosystems, Carlsbad, CA). All samples were run in duplicate. For comparison purposes, mRNA from sham rats were always run within the same plate as experimental samples from vehicle treated and antalarmin treated groups.

### Statistical analyses

GraphPad Prism 6.0 (Graph-Pad Software, San Diego, California) was used to prepare graphs and run statistical analyses. Data is presented as mean difference ± SEM and a p value <0.05 was considered statistically significant. The variability between groups was first assessed followed by a test for outlier values. A Student t-test was used for comparisons between two groups and when group variability was significantly different, a Welch corrected t-test was used. A one-way analysis of variance (ANOVA) was used to compare behaviors. A one-sample t-test against the sham rats value normalized to 1.0 was used to assess qRT-PCR results. A repeated measures one-way ANOVA was used to compare changes in weight gain between treatment groups across time.

## Results

### Shortly after endometriosis induction, CRHR1 is elevated

To evaluate the early endometriotic vesicle development, we induced the disease and sacrificed the animals after 7 days. It is during this early development period that we hypothesized that the major changes in CRHR1 will be observed. At 7 days, we observed that 93.7 % of the implants have created a large vesicle, which in most cases was very large and filled with fluid. In the sham group, only sutures were observed as no uterus was transplanted. Table 1 shows the morphological characteristics of the observed vesicles at 7 days post-induction surgery. We quantified the CRHR1 mRNA in the vesicles as compared to the normal uteri of the same rats and that of sham surgery controls. CRHR1 mRNA in endometriosis vesicles showed a two-fold increase as compared to normal uterus of sham rats (t=2.934, d.f.=6, p<0.05; Figure 2). In contrast, the mRNA levels in uteri of rats that received endometriosis were not different from the uteri of sham rats (t=0.829, d.f.=6, p>0.05; Figure 2).

**Figure 2:**
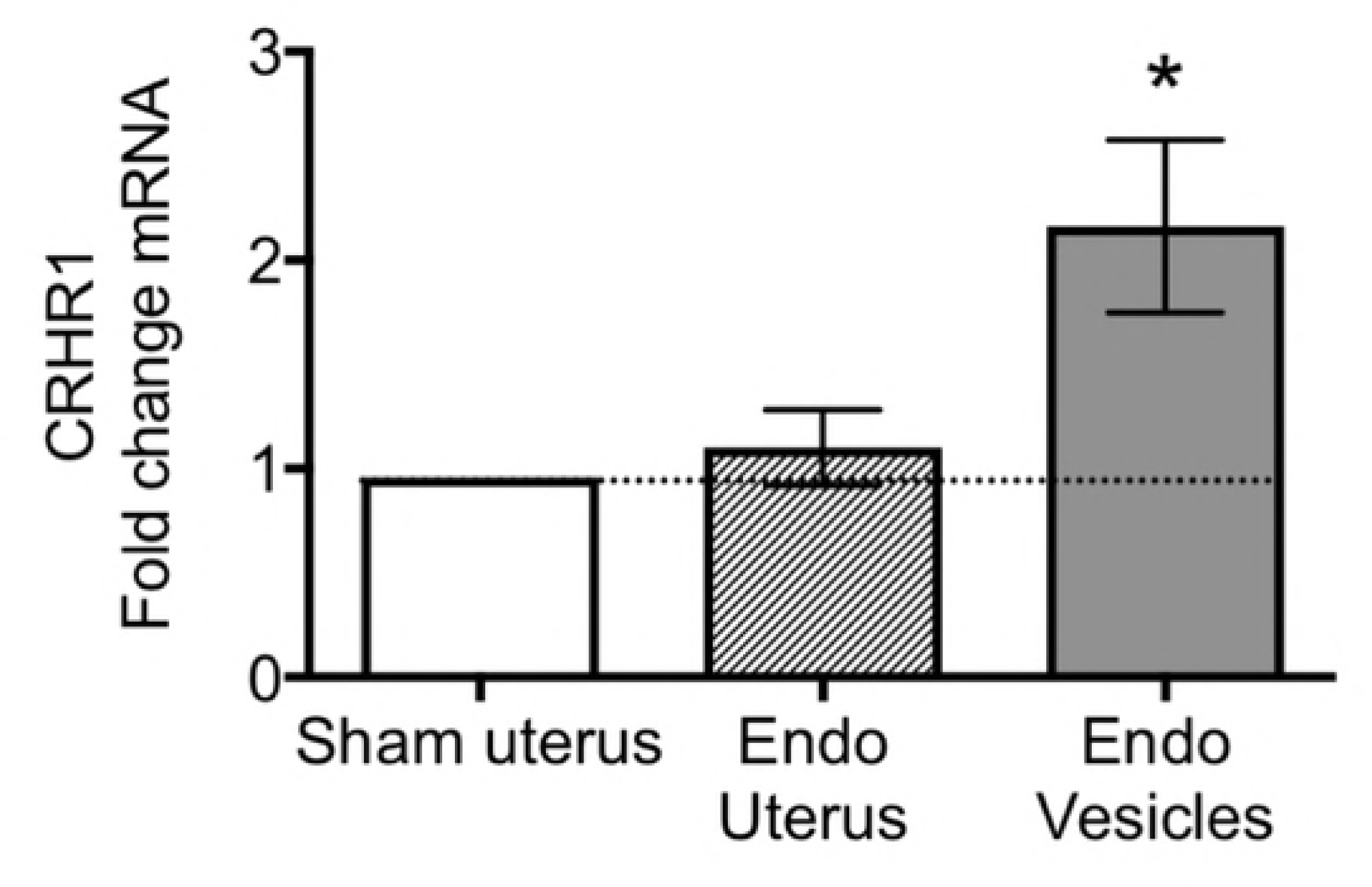
qRT-PCR of CRHR1 within endometriosis vesicles and uterus of the endometriosis rats and sham rats. At 7 days after the autotransplantation surgery to induce endometriosis, we observed a significant two-fold increase in CRHR1 mRNA within endometriosis vesicles only.

**Table 1:** Characteristics of endometriosis vesicles at seven days after auto-transplantation surgery.

| Endo vesicles at 7 days | Percent developed (%) | Total weight (g) | Total area (mm<sup>2</sup>) | Total volume (mm<sup>3</sup>) |
| --- | --- | --- | --- | --- |
| Average per rat ± S.E.M. | 93.75 ± 2.80 | 1.19 ± 0.25 | 160.92 ± 15.64 | 2190.25 ± 1110.58 |

### Antalarmin did not affect stress reactivity or anxiety behaviors

To block the significant increase in CRHR1 receptor within the endometriotic vesicles, we administered antalarmin during the first 7 days after endometriosis induction surgery. After that, we allowed the endometriosis to progress for 53 additional days. At Day 59 after endometriosis surgery, a subset of the animals (6 vehicle and 6 antalarmin) were subjected to a 5-min swim stress challenge. Antalarmin treated animals were not different from the vehicle control group in any of the behavioral parameters measured such as immobility, swimming, struggling behaviors and diving episodes (data not shown). Therefore, rats tested in the stress challenge were collapsed within the not-tested ones within treatment groups (vehicle or antalarmin). The next day, all animals were tested using the open field and the zero maze to evaluate trait and state anxiety, respectively. The total distance traveled (Figure 3A) was significantly lower in rats that received antalarmin as compared to the sham group (F_(2,45)_= 3.507, p< 0.05), but the amount of time rats spend in the center of the open field arena was similar between groups (F_(2,45)_= 0.12, p> 0.05; Fig. 3B). On the zero maze, a higher locomotor activity was observed for both groups of rats with endometriosis that received vehicle or antalarmin compared to the sham group (F_(2,49)_= 6.01, p< 0.01; post-hoc, p≤ 0.01 both comparisons; Fig. 3C). Despite a higher locomotor activity, there was a strong trend for both groups of rats with endometriosis to spend less time in the open arms of the zero maze ( F_(2,49)_= 2.82, p= 0.06; Fig. 3D) suggesting increased anxiety. In summary, antalarmin administered shortly after endometriosis induction does not have long-term effects on anxiety behaviors. However, endometriosis tended to increased anxiety in the zero-maze compared to sham controls, regardless of treatment.

**Figure 3:**
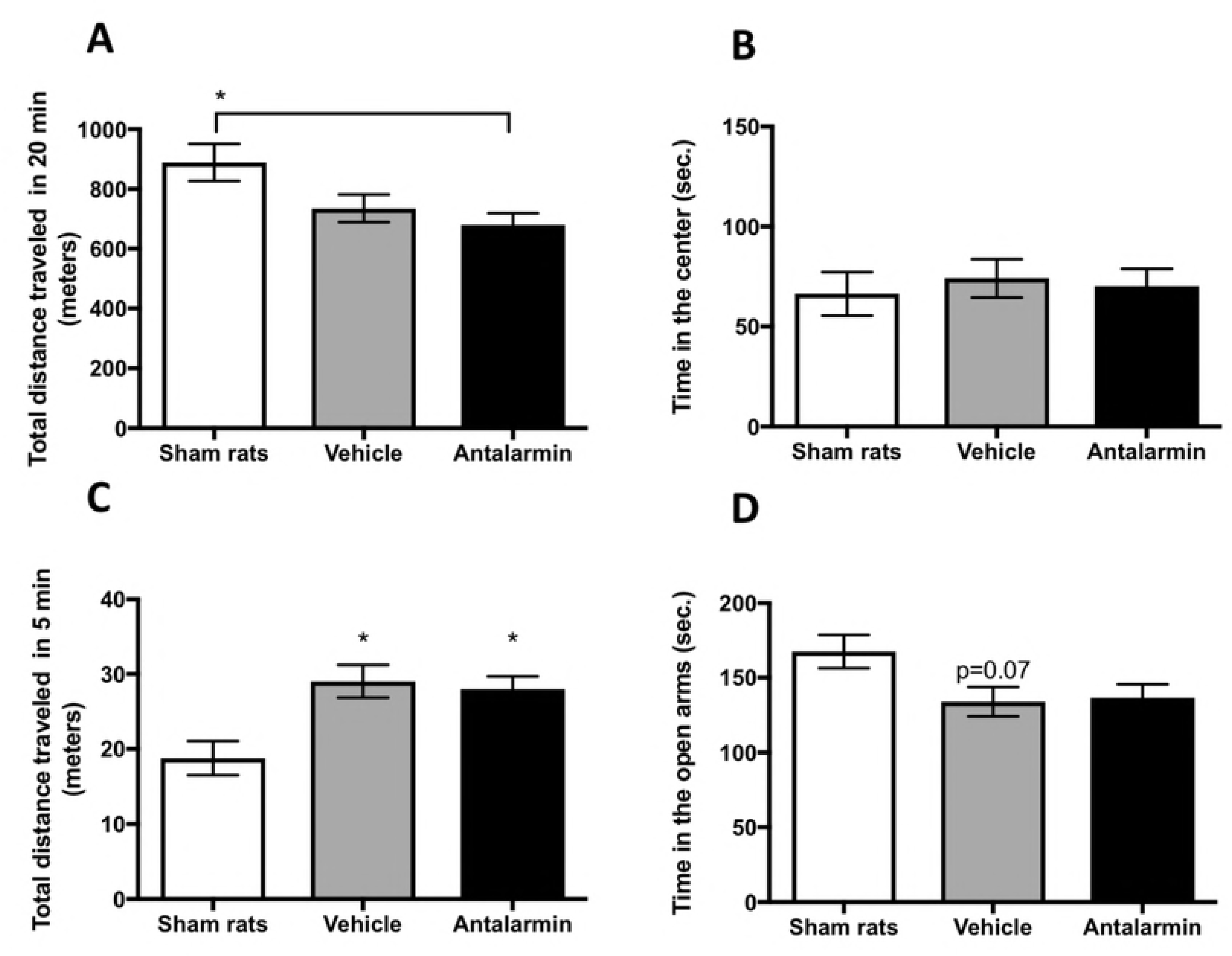
Behavioral assessment for anxiety. Rats that received sham surgery or endometriosis and antalarmin or vehicle treatment were tested in the open field (A and B) or the elevated zero maze (C and D). In comparison to sham, we observed a significant decrease in locomotor activity of the group that received antalarmin. However, time spent in the center of the open field was not different between groups. In the elevated zero maze, a significant increase in locomotion was observed for rats that had endometriosis as compared to sham, regardless of the drug treatment (C). Rats with endometriosis, regardless of drug treatment, showed a trend towards spending less time in the open segment of the zero maze as compared to sham group. * represents p< 0.05.

### Antagonizing CRHR1 early in endometriosis produced a significant decrease in vesicle development

Antalarmin administration during the 7 days after endometriosis induction resulted in a 30% significant decrease in the number of developed endometriosis vesicles at 60 days (Welch corrected t-test, t= 3.38, d.f.= 22.86, p<0.01; Fig. 4A). The total weight of endometriosis vesicles (sum per rat) in the antalarmin treated group was 67% less than the vehicle control group (t= 3.175, d.f.= 38, p<0.01; Fig. 4B). The reduced weight was a direct result of the smaller size of the vesicles in average volume (68% difference, Welch corrected, t= 2.515, d.f.= 25.39, p<0.05; Fig. 4C) and area (55% difference, t= 3.067, d.f.= 38, p<0.01; Fig. 4D) per rat. Similar to our previous reports, (21,30) we classified the vesicles in grades (1 – 5) based on a length scale for each vesicle where 1 denotes an implant that disappeared and 5 an implant that developed into a vesicle equal or larger than 6mm (Fig. 4E). In the antalarmin treated group compared to the vehicle treated control, there was a larger percentage of endometriosis vesicles that disappeared, as well as a reduced percentage of vesicles of grade 3 and 5. In summary, seven days of antalarmin treatment resulted in a smaller percentage of endometriosis implants developed and those that did develop were significantly smaller in size compared to vehicle treated control group.

**Figure 4:**
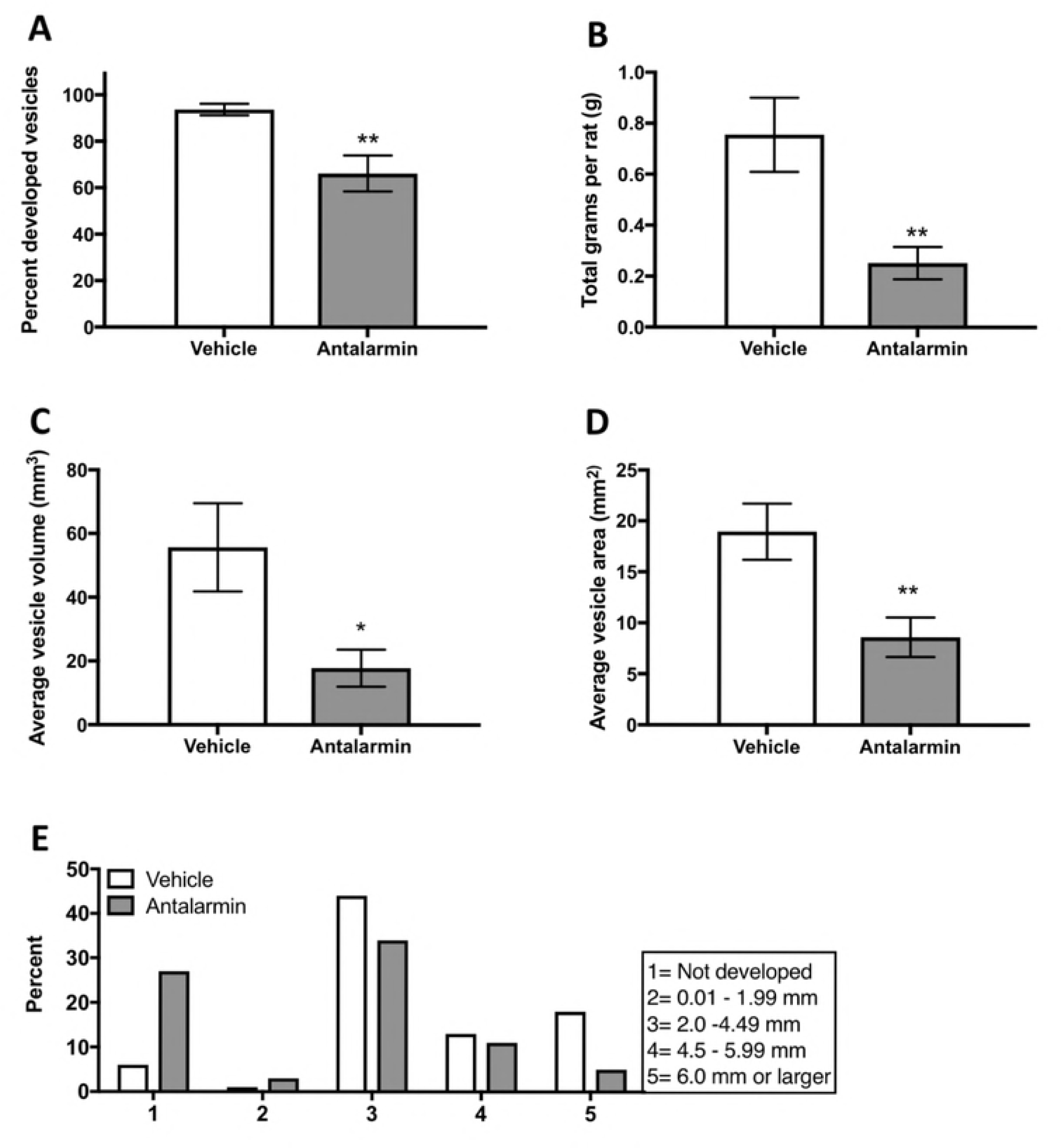
Morphological characteristics of endometriosis vesicles. (A) The percent of implants that developed into vesicles was significantly lower in the antalarmin treated group compared to the vehicle control group. (B) The total weight of all vesicles per rat was smaller for the antalarmin treated rats. (C) The average vesicle volume per rat was significantly smaller for the antalarmin treated group compared to the vehicle control group. (D) The average vesicle area per rat was significantly smaller in the antalarmin group compared to the vehicle group. (E) Vesicles that developed were classified by grade based on a scale by size. * p< 0.05, ** p< 0.01.

### Antalarmin produced a long-lasting increase in serum ACTH

At the time of sacrifice, we collected peritoneal fluid and blood serum from rats to later examine how the treatment with antalarmin might have altered HPA axis markers and also the pro-inflammatory cytokine IL-6. Corticosterone was slightly elevated in rats with endometriosis treated with vehicle or with antalarmin, however this apparent difference, did not reach statistical significance (F_(2,43)_= 1.99, p>0.05; Fig. 5A). On the other hand, we observed significantly elevated levels of serum adrenocorticotropic hormone (ACTH) in rats that received antalarmin. ANOVA statistical test revealed a significant main effect of drug treatment (F_(2,39)_= 518, p<0.05; Fig. 5B). Post hoc tests showed that the groups of rats that received antalarmin was significantly higher that sham and vehicle treated groups (p< 0.05 both comparisons). While anti-inflammatory effects of antalarmin have been reported, the short treatment of antalarmin 53 days before sacrifice did not produce any change in IL-6 in peritoneal fluid, which is in direct contact with the endometriotic vesicles (Fig. 5C).

**Figure 5:**
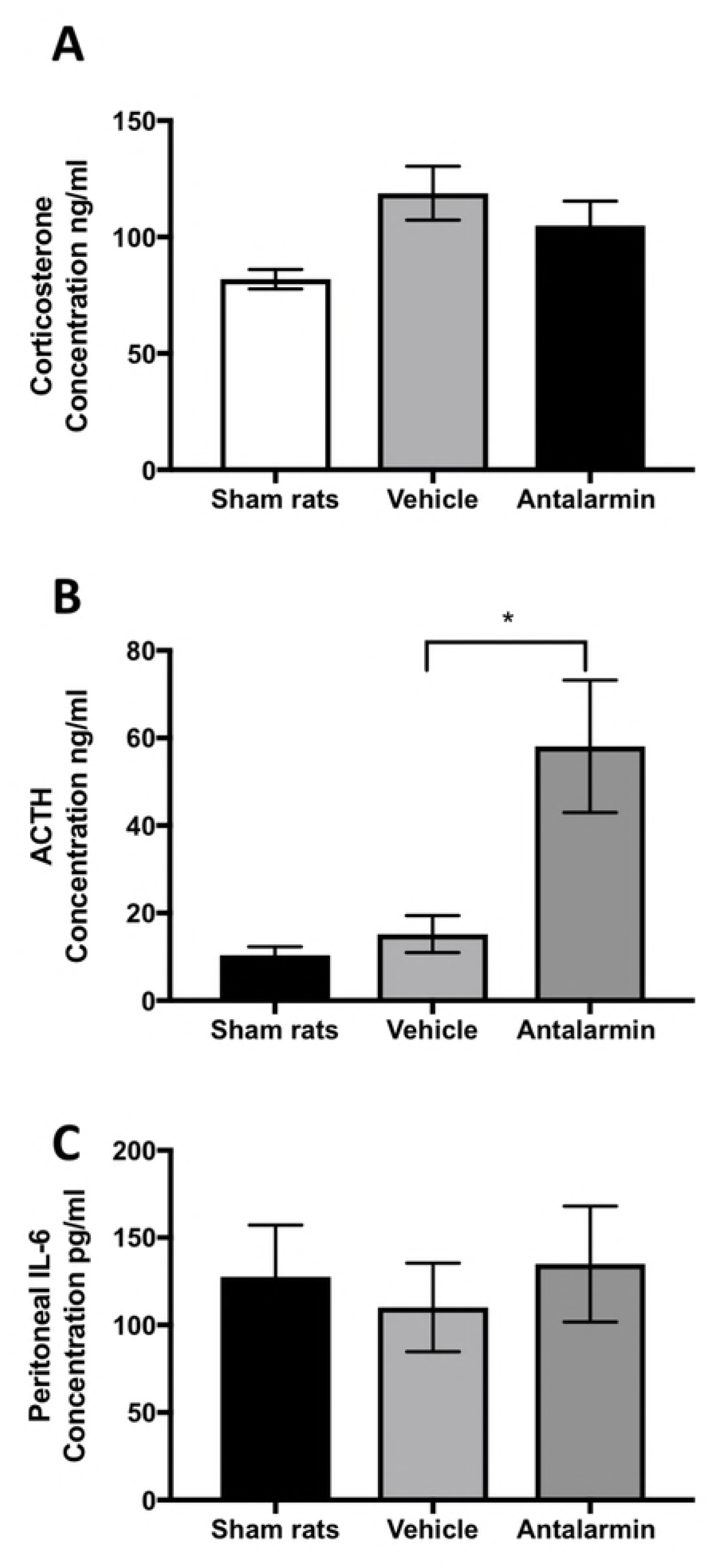
Serum and peritoneal markers of stress and inflammation. We used ELISA to measure (A) corticosterone and (B) adrenocorticotropic hormone (ACTH) in the serum of rats and (C) IL-6 in peritoneal fluid at the time of sacrifice. There was no significant difference between groups in serum corticosterone levels at the time of sacrifice. However, a significantly higher level of ACTH was observed for rats that received antalarmin compared to the two other groups. No differences in IL-6 were observed between groups. * represents p< 0.05.

### Antalarmin blocked mRNA increase in CRH and CRHR1 of uterus and vesicles

We quantified the mRNA for urocortin and CRH, which are the main agonists of the CRHR1 receptor, within developed endometriosis vesicles in rats from both treatment groups using qRT-PCR. As a comparative parameter, we also quantified the mRNA with the uteri of the same animals and used uteri of sham controls to normalize the data. We observed a significant two-fold increase in CRH for vehicle treated rats, both in uterus (one sample t-test: t= 2.66, d.f.= 13, p< 0.05) and vesicles (t= 2.29, d.f.= 13, p< 0.05; Fig. 6A). However, this increase was not observed in the antalarmin treatment group (Fig. 6A). In contrast, UCN1 mRNA was not altered in any of the groups measured (Fig. 6B).

**Figure 6:**
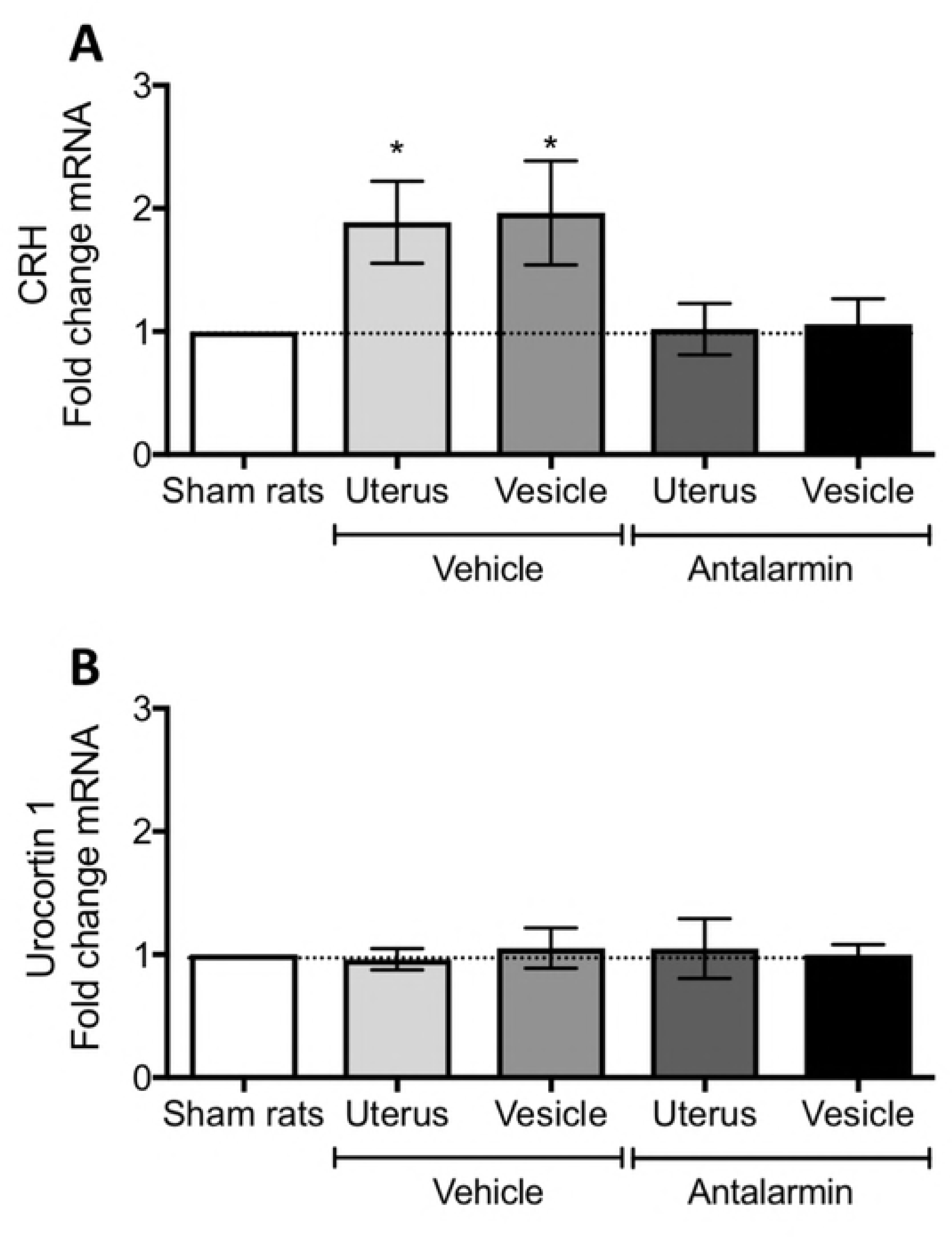
mRNA levels measured by qRT-PCR from the uterus and endometriosis vesicles. (A) corticotropin releasing hormone (CRH). (B) Urocortin 1 peptide. Data normalized to the uterus of sham rats. * represents p< 0.05 compared to sham rats’ uterus.

The mRNA of the CRHR1 receptor measured in endometriosis vesicles of the vehicle group was significantly increased as compared to sham uterus (t= 3.45, d.f.= 8, p< 0.01; Fig. 7A), but this increase was not observed in the vesicle of antalarmin treated rats (p>0.05). Due to the intricate balance of CRH receptor activity in uterine tissue, we also quantified the CRHR2 receptor mRNA. For this receptor, we observed a small but significant fold increase in mRNA only in the vesicles of vehicle treated animals (t= 3.2, d.f.= 8, p< 0.05; Fig. 7B). No other changes were observed for CRHR2. The glucocorticoid receptor showed an interesting pattern with a significant mRNA fold increase that was observed in vesicles from both, the vehicle treated (t= 2.88, d.f.= 8, p< 0.05; Fig. 7C) and antalarmin treated (t= 4.65, d.f.= 8, p< 0.01; Fig. 7C) groups, but no changes in uterus.

**Figure 7:**
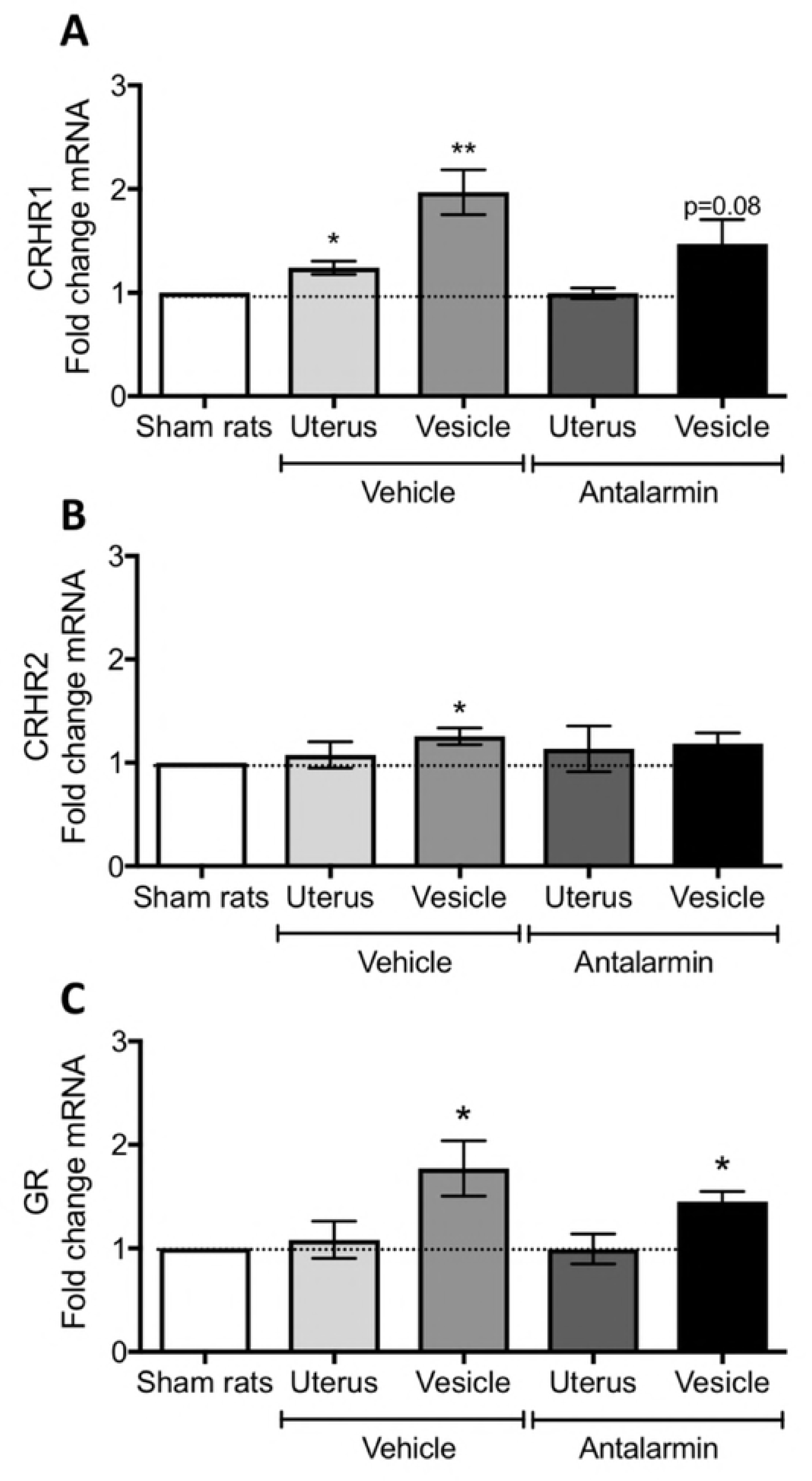
mRNA levels measured by qRT-PCR from the uterus and endometriosis vesicles. (A) corticotropin releasing hormone receptor type 1 (CRHR1). (B) corticotropin releasing hormone receptor type 2 (CRHR2). (C) glucocorticoid receptor (GR). Data normalized to the uterus of sham rats. * represents p< 0.05 compared to sham rats’ uterus.

### Antalarmin decreased body weight and the decrease persisted in treated animals

Inconsistencies in the effect of antalarmin on male rodent body weight have been reported (34,35). We monitored the rat weight changes during the drug administration period and every week afterwards during the development of endometriosis. By day 7 and for 2 days after antalarmin has stopped, rats receiving the drug weighed significantly less than vehicle control groups (Repeated measures ANOVA of treatment: F_(1,38)_= 7.615, p< 0.01, post hoc, p<0.01 on day 7 and p< 0.05 on days 8 and 9; Fig. 8A). While the rats maintained a constant weight gain rate, by week 7 and 8 after endometriosis induction (Figure 8B), a significantly lower weight gain compared to vehicle group was also recorded (F_(1,38)_= 33.89, p< 0.001, post hoc, p<0.05 for weeks 6 and 7).

**Figure 8:**
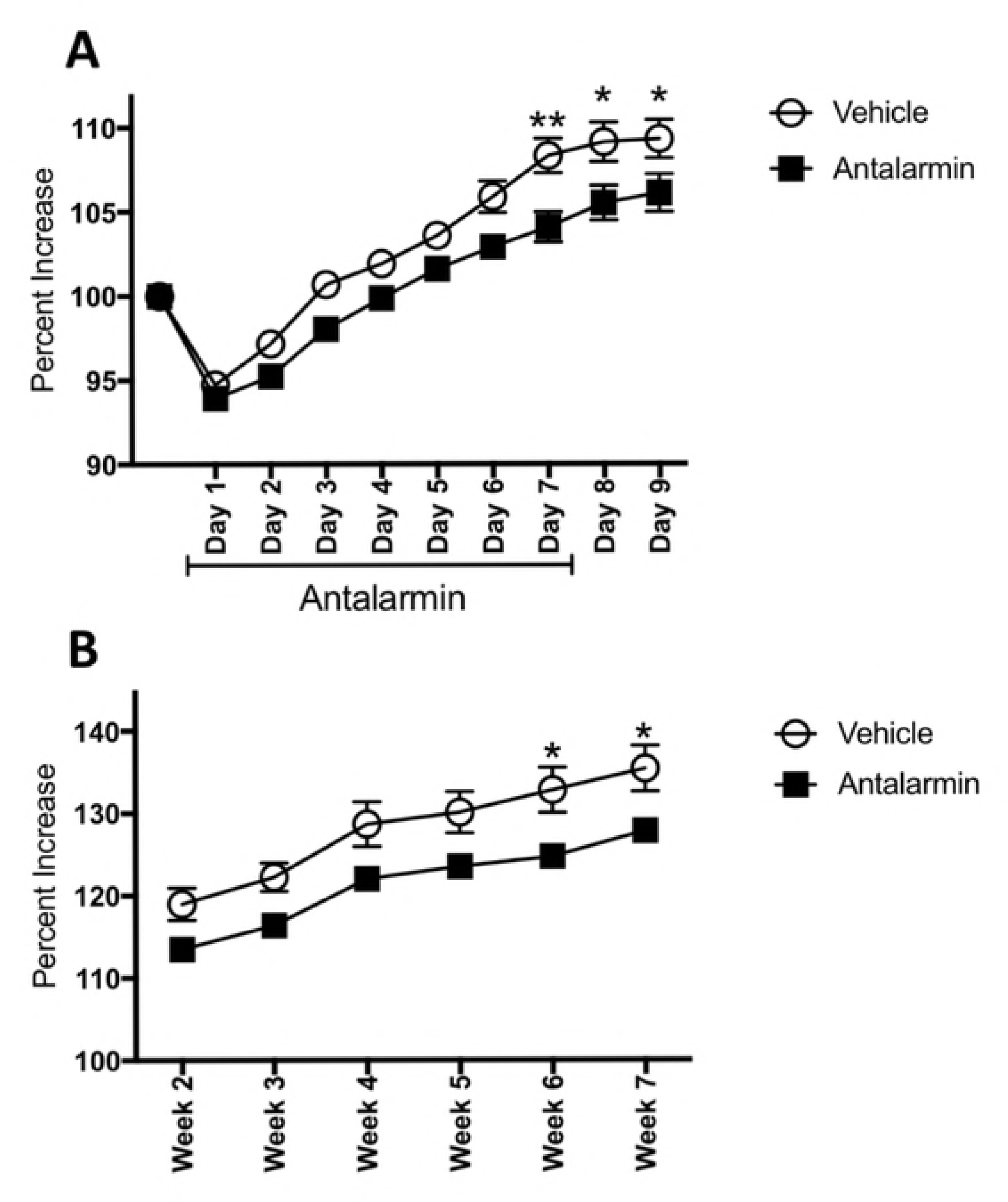
Percent increase in weight for rats with endometriosis treated with antalarmin or vehicle. (A) After seven days of treatment, antalarmin treatment significantly decreased the weight of the rats and this difference persisted for two additional days after the drug treatment has stopped. (B) During the subsequent weeks, antalarmin group remained weighing less than control group and by weeks 6 and 7 this difference reached statistical significance. * p< 0.05, ** p< 0.01.

## Discussion

Here we present evidence that a short treatment with the CRHR1 antagonist antalarmin had long-term efficacy in reducing endometriosis with minimal changes in behavior. More importantly, the vesicles that developed were significantly smaller, suggesting that antalarmin interfered with both the establishment and the development of vesicles. We also demonstrated that antalarmin prevented the increase in CRH and CRHR1 mRNA within endometriosis vesicles as compared to vehicle treated rats that lasted for almost 2 months after treatment stopped. This suggests that a short treatment might produce long-lasting effects for the treatment of this disease. To our knowledge, our work provides the first *in vivo* evidence of efficacy and use of CRHR1 antagonist antalarmin for the treatment of endometriosis.

In the present study, only one time point was tested corresponding to a major increase in CRHR1 mRNA, early in endometriosis development, as demonstrated herein. Sampson’s theory postulates that retrograde menstruation leads to endometriosis. Based on this theory, during every menstruation, there is opportunity for new endometriosis implants to develop. Therefore, we suggest that in the clinical scenario, treatment with CRHR1 antagonist would need to be used right after laparoscopic surgery or for several years, similar to contraceptive pills, in order to be effective. In our animal model, we observed that about 60% of the implants sites developed into endometriosis vesicles. Perhaps a longer or continuous treatment with the CRHR1 antagonist might have produced a larger decrease in endometriosis vesicle development or even completely abolish it. In the clinical setting, there is a significant lag in the diagnosis of endometriosis, which is on average 7 years after symptoms appear. Our work opens the possibility for future testing of antalarmin or other CRHR1 antagonist at later time points in disease progression.

Not all CRHR1 antagonists are equal. The clinical use of CRHR1 antagonists has been limited by several factors that include lack of consistent efficacy (36,37), elevated tissue accumulation and prolonged half-life (38,39). Recently, a group of orally administered CRHR1 antagonists have been shown to have high bioavailability and low lipophilicity in animal models of IBS (40). The availability of these new antagonists opens significant possibilities for the advancement of testing new CRHR1 compounds in endometriosis. However, certain challenges still remain. Eleven isoforms of the CRHR1 receptor have been identified in humans (8), and splicing of CRHR1 seems to be tissue specific. For example, CRHR1β is present in pituitary myometrium and endometrium but not in adrenal, placenta or synovium (41,42). It still needs to be determined whether ectopic endometrium will display a different profile of CRHR1 splice variants as compared to eutopic endometrium both in pre-clinical studies as well as in the clinical setting.

One of the most interesting findings of the current study was the elevated levels of ACTH in plasma that persisted even after antalarmin administration stopped. However, this elevated response was not followed by an increase in plasma corticosterone (see Fig.5). ACTH binds to the melanocortin receptor type 2 (MC2) to stimulate the release of glucocorticoids from the adrenal glands (43). In human endometrium, all five types of melanocortin receptors have been found (MC1-5) and when exposed to ACTH, decreased vascularity was observed in cultured decidual biopsies (44). Therefore, some of the long-term effects of antalarmin on decreasing endometriosis progression in our current study might be attributed to an increased level of circulating ACTH, resulting in decreased endometrial vesicle development. While further experiments are necessary to elucidate the increase in ACTH without a concomitant increase in corticosterone observed herein, the most plausible explanation is a de-sensitization of intra-adrenal signaling system. In support of this, there is evidence that human adult adrenal tissues express, ACTH, CRH, CRHR1 and CRHR2 mRNA and that exposure of adrenal cells to antalarmin blocks the production of cortisol (45). Therefore, CRHR1 are involved in the control of glucocorticoid secretion, and antalarmin administration might have led to long-term dampening or desensitization within adrenal tissues.

Long-term changes in behavior due to the antalarmin administration were minimal. This suggest that compensatory activity most likely occurred in the amygdala and/or other regions involved in controlling anxiety-like behaviors. A recent study using intracerebroventricular administration of antalarmin showed that blocking CRHR1 provides neuroprotection and blunts neuroinflammation resulting from global cerebral ischemia (46). Blocking CRHR1 in the hippocampus results in a reduction of excitatory activity onto CA3 pyramidal cells in hippocampus (47). Clinically, CRHR1 signaling has been implicated in mediating abnormal brain responses to expected abdominal pain in patients with IBS (48). Based on the significant role of CRHR1 in homeostasis and behaviors, one of the challenges is to optimize CRHR1 peripheral blockade, while producing minimal changes in the brain. This provides an opportunity for drug design that targets peripheral blockade of CRHR1 but prevents lasting effects in brain and behavior.

One of the challenges in the clinical setting is to decrease endometriosis sites, while still preserving reproductive abilities. Antalarmin has been shown in rodents to reduce the number of implantation sites by 70% (49) by a Fas-ligand immune tolerance dependent mechanism (50). It is possible that antalarmin treatment will compromise reproductive abilities. However, the long-lasting effect of antalarmin in the current study opens the possibility of developing short-term treatments that will provide long-lasting protection and allow for reproductive abilities to return to normal. This still remains to be tested.

## Conclusion

A single week of antalarmin treatment was effective in reducing endometriosis in the rat model by reducing the number of developed vesicles by 30 % and the size of the vesicles that developed by 67%. CRHR1 Inhibitors are pharmacological agents that are advanced in the pipeline of clinical trials in safety and efficacy profiles for other inflammatory disorders such as IBS. Our study opens the possibility for a new application of CRHR1 inhibitors for the treatment of endometriosis. We predict that translation of our work into the clinical application can produce significant benefits for many women that suffer from endometriosis.

## Acknowledgements

This work was funded by the following NIH grants: K07AT008027 to A.T.R. and the RCMI BRAIN (behavioral) and MAGIC (molecular) cores MD007579 and by RISE grant 2R25GM082406. The authors acknowledge the participation of graduate students Inevy Seguinot and Raura Doreste from Ponce Health Sciences University and undergraduate student Yammile Vargas from University of Texas at Rio Grande Valley. We would like to acknowledge the technical assistance of Maria del C. Colon in the behavioral experiments. The helpful comments of Dr. Siomara Hernandez and Dr. Bruce McEwen are also greatly appreciated.

